# Spatial simulation of co-designed land-cover change scenarios in New England: Alternative futures and their consequences for conservation priorities

**DOI:** 10.1101/722496

**Authors:** Jonathan R. Thompson, Joshua Plisinski, Kathy Fallon Lambert, Matthew J. Duveneck, Luca Morreale, Marissa McBride, Meghan Graham MacLean, Marissa Weis, Lucy Lee

## Abstract

To help prepare for an uncertain future, planners and scientists often engage with stakeholders to co-design alternative scenarios of land-use change. Methods to translate the resulting qualitative scenarios into quantitative simulations that characterize the future landscape condition are needed to understand consequences of the scenarios while maintaining the legitimacy of the process. We use the New England Landscape Futures (NELF) project as a case study to demonstrate a transparent method for translating participatory scenarios to simulations of Land-Use and Land-Cover (LULC) change and for understanding the major drivers of land-use change and diversity of plausible scenarios and the consequences of alternative land-use pathways for conservation priorities. The NELF project co-designed four narrative scenarios that contrast with a *Recent Trends* scenario that projects a continuation of observed changes across the 18-million-hectare region during the past 20 years. Here, we (1) describe the process and utility of translating qualitative scenarios into spatial simulations using a dynamic cellular land change model; (2) evaluate the outcomes of the scenarios in terms of the differences in the LULC configuration relative to the *Recent Trends* scenario and to each other; (3) compare the fate of forests within key areas of concern to the stakeholders; and (4) describe how a user-inspired outreach tool was developed to make the simulations and analyses accessible to diverse users. The four alternative scenarios populate a quadrant of future conditions that crosses high to low natural resource planning and innovation with local to global socio-economic connectedness. The associated simulations are strongly divergent in terms of the amount of LULC change and the spatial pattern of change. Features of the simulations can be linked back to the original storylines. Among the scenarios there is a fivefold difference in the amount of high-density development, and a twofold difference in the amount of protected land. Overall, the rate of LULC change has a greater influence on forestlands of concern to the stakeholders than does the spatial configuration. The simulated scenarios have been integrated into an online mapping tool that was designed via a user-engagement process to meet the needs of diverse stakeholders who are interested the future of the land and in using future scenarios to guide land use planning and conservation priorities.

## INTRODUCTION

Scenario planning is a rigorous way of asking “*what if?*” and it can be a powerful tool for natural resource professionals preparing for the future of socio-ecological systems. In the context of land-use or regional planning, scenario development uses a structured process to integrate diverse modes of knowledge to create a shared understanding of how the future may unfold (MA 2005, Mahmoud et al. 2009, Wiebe et al. 2018). The resulting scenario narratives that emerge from participatory scenario planning describe alternative trajectories of landscape change that would logically emerge from different sets of assumptions (Thompson et al. 2012). Scenarios are not forecasts or predictions; instead, they are a way to explore multiple hypothetical futures in a way that recognizes the irreducible uncertainty and unpredictability of complex systems (Pedde et al. 2018).

Scientists are increasingly co-designing scenarios with stakeholders—i.e., groups of people who are both affected by and/or can affect decisions or outcomes (Voinov and Bousquet 2010, Reed et al. 2013, McBride et al. 2017). Co-designing scenarios increases the range of viewpoints and expertise included in the process and, in turn, attempts to increase the relevance, credibility and salience of outcomes (sensu, Cash et al. 2003). Participatory land use scenario development is particularly useful in landscapes such as New England where landscape change is driven by the behaviors and decisions of hundreds of thousands of stakeholders that are not amenable to centralized planning or prediction. A land-use scenario co-design process typically results in a set of contrasting storylines that describe the way the future might unfold, based on specific assumptions about dominant social and ecological forces of change within a landscape (Ramírez and Selin 2014, McBride et al. 2017).

The utility of qualitative, co-designed scenarios can be enhanced by linking them to quantitative representations of future land-use change, as generated by a spatially explicit simulation model. However, translating between narrative scenario descriptions and quantitative models presents challenges and tradeoffs related to the treatment of uncertainty, the potential to accommodate stakeholders in the process, the resources required, and the compatibility with different types of simulation models (see reviews of these factors in: Mallampalli et al. 2016, Pedde et al. 2018). These challenges notwithstanding, variations on the “Story and Simulation” approach (sensu Alcamo 2008) to scenario planning are increasingly used in in environmental planning and are the basis for many large-scale regional scenario assessments (MA 2005, Rounsevell et al. 2006, Thompson et al. 2014, 2016, Carpenter et al. 2015, Sohl et al. 2016, Kline et al. 2017).

Cellular land change models (LCM) have features that make them well suited to the translation of qualitative scenarios to spatial simulations (Brown et al. 2013, Dorning et al. 2015, Thompson et al. 2017). Cellular LCMs are phenomenologically driven, as opposed to process-driven, and are often used to project observed trends of land use and land cover (LULC) change forward in time. By projecting observed trends of LULC change, they operate with the implicit assumption that the future will be a continuation of the past (e.g., Thompson et al. 2017). These models quantify the rate of LULC change and the relationships between the location of observed LULC change (i.e., a change detection) and a suite of spatial predictor variables--e.g., patterns of existing development, proximity to city centers or roads, topography, demographics etc. Simulating these patterns into the future constitutes a “recent trends” scenario, which can be used as a baseline, against which alternative scenarios can be evaluated. Then, by adjusting LULC change rates and/or re-defining the strength or nature of the relationships between LULC changes and spatial predictor variables, modelers can systematically and transparently simulate alternative scenarios. Cellular LCMs can also incorporate feedbacks to LULC change and portray multiple interacting land uses. For example, on a simulated forested site, new land protection can prevent new residential development from occurring. New residential development in a simulation can also increase the probability that additional new development will occur in proximity to existing development. This dynamic modelling approach produces a realistic manifestation of LULC change by re-producing observed landscape patterns (Wilson et al. 2003). Finally, cellular LCMs are relatively straightforward to understand and to describe to stakeholders, as compared with agent-based or other more computationally sophisticated approaches to land-use simulation (Brown et al 2013).

### New England Landscape Futures

Here we use the New England Landscape Futures (NELF) project as a case study to demonstrate the potential for translating participatory scenarios to simulations of LULC change and for understanding the consequences of alternative land-use pathways for conservation priorities. NELF is a multi-institutional, participatory scenario project with the overarching goal of building and evaluating scenarios that show how land-use choices and climate change could shape the landscape over the next 50 years.

The six-state, 18-million-hectare region has several characteristics that lend itself to participatory scenario planning (McBride et al. 2017). Seventy-five percent of New England forests are privately owned, including the nation’s largest contiguous block of private commercial forestland (> 4 million ha) plus hundreds of thousands of family forest owners with small to mid-sized parcels totaling > 7 million hectares (Butler et al. 2016). It is among the most forested and most populated regions in the U.S.; average forest cover in the region exceeds 80% but ranges from 50% in Rhode Island to 90% in Maine (Figure 1). The future of these forests is in question. Since 1985, roughly 10,000 ha yr-1 of forest have been lost to commercial, residential, and energy development, marking the reversal of a 150-year period of forest expansion in the region (Olofsson et al. 2016). Working to slow the rate of forest loss are a range of robust conservation initiatives that have, to date, permanently protected 23% of the region from development; half of this conservation land has been protected since 1990 (Foster et al. 2017, Sims et al. 2019). Modern land protection in this region is primarily achieved by private land owners voluntarily placing conservation restrictions on their land. Likewise, development of forest or agricultural sites to residential or commercial uses is made primarily by individual private land owners. Thus, these individual choices are collectively determining the future of the shared landscape. There is no central decision-making authority for land use; instead, the condition of future landscape will be the product of countless independent landowner decisions and a conglomerate of local, regional, and state policies.

**Figure 1.**
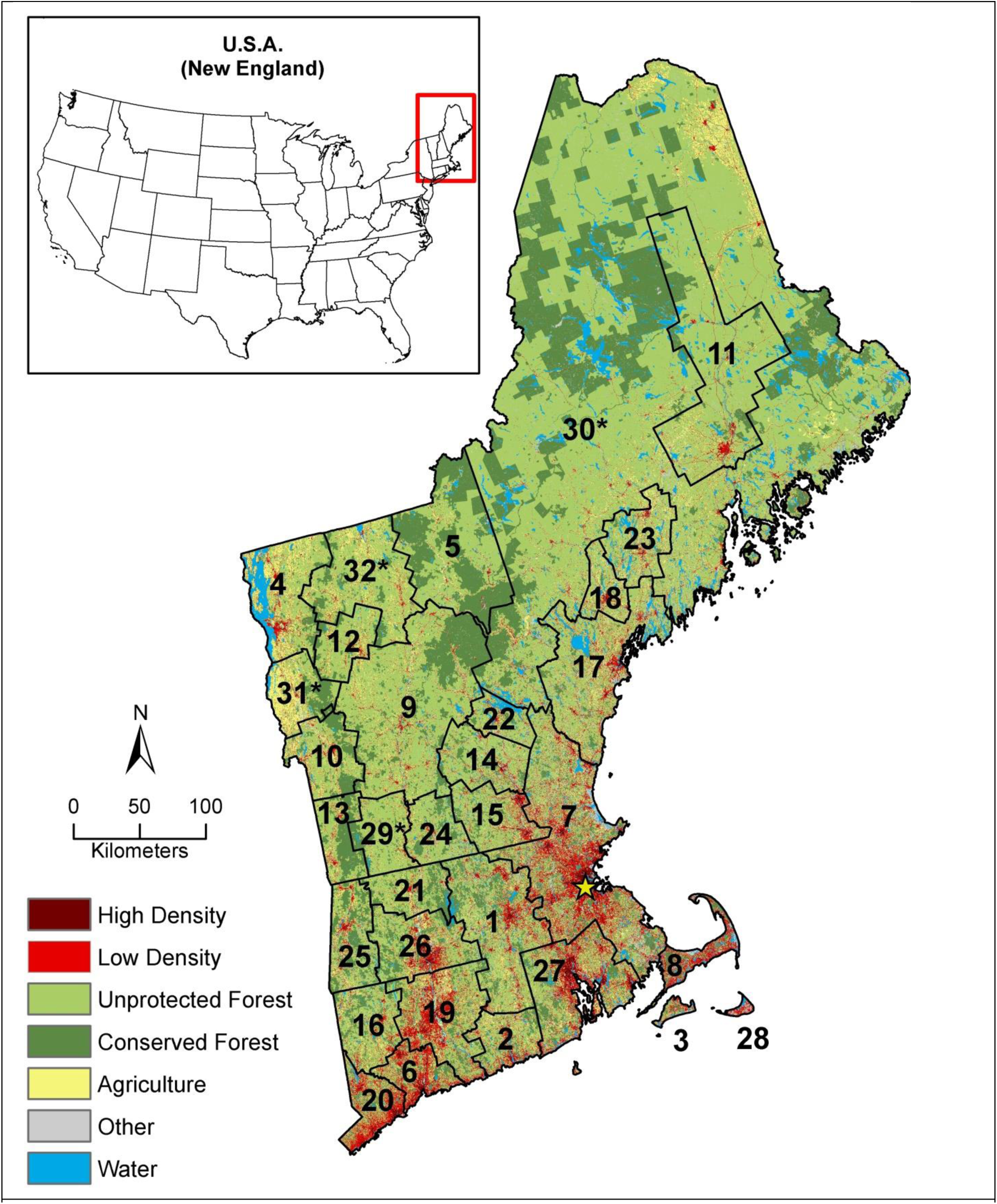
Study Area with numbered subregions. Asterisk denotes non-CBSA subregions.

McBride et al. (2017) describe the participatory process through which the NELF project codesigned four divergent narrative scenarios that contrast with a *Recent Trends* scenario. In brief, four scenarios were co-designed through a structured scenario development process that engaged > 150 stakeholders and scientists from throughout the study region. Using the Intuitive Logics approach to scenario development popularized by Royal Dutch Shell/Global Business Network (Bradfield et al. 2005), the NELF project stakeholders envisioned opposing outcomes of two key drivers of land-use change that they identified as highly impactful and highly uncertain: socio-economic connectedness and natural resource planning and innovation. The process resulted in a matrix of four quadrants that encompassed four broad scenarios. Participants then added details about each scenario storyline in qualitative terms, which took the form of ∼1000 word narratives (McBride et al. 2017) and are summarized in the Scenario Narratives (Table 1). Next, participants were presented with key features of the *Recent Trends* scenario and asked to describe how land use would differ in each of the alternative scenarios using semi-quantitative terms. We then adjusted model input parameters to reflect the characteristics of each of the four divergent scenarios. Finally, through a series of subsequent interactive webinars we worked with participants to refine these parameters to ensure the scenarios captured their intent.

**Table 1.**
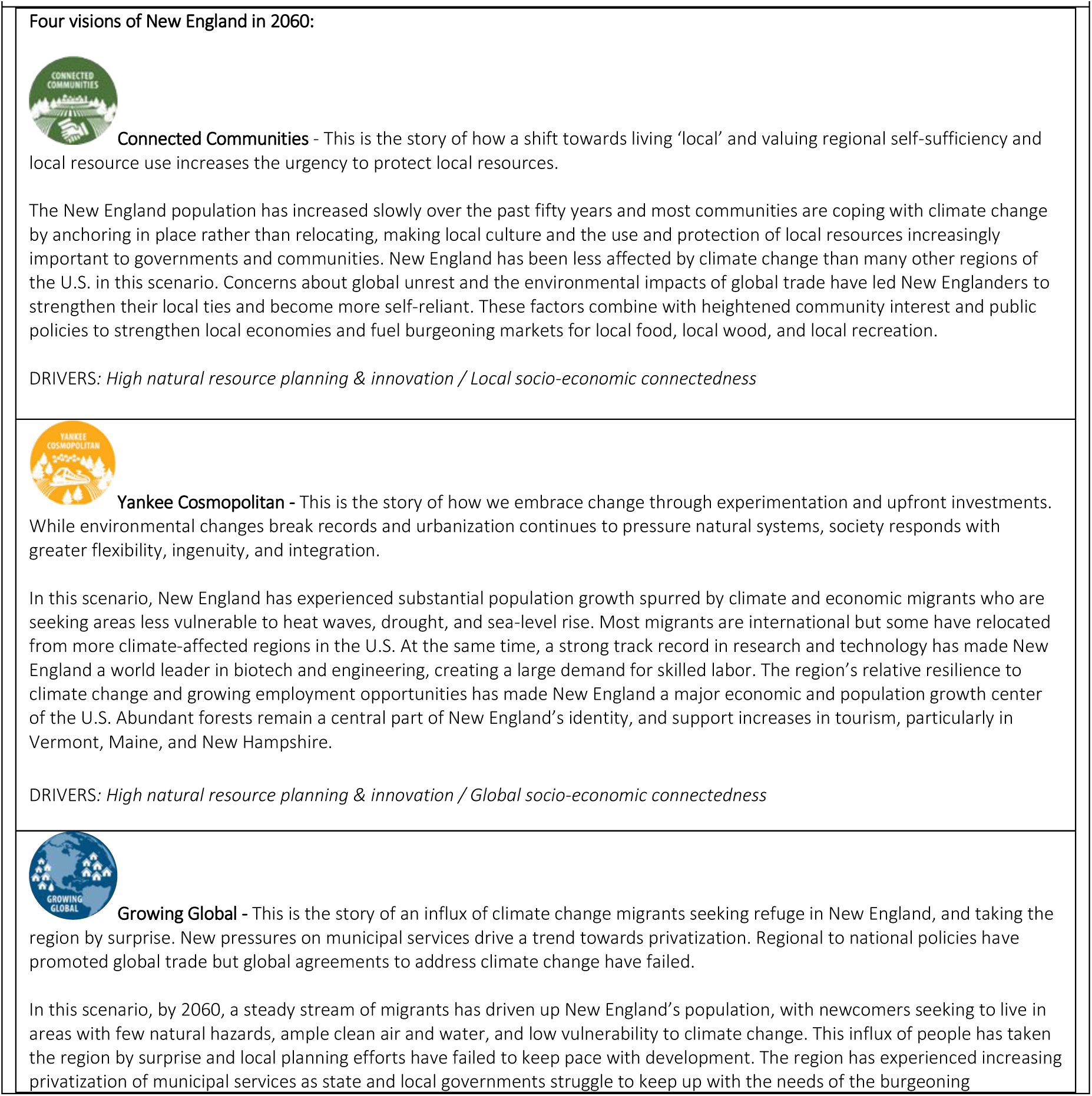
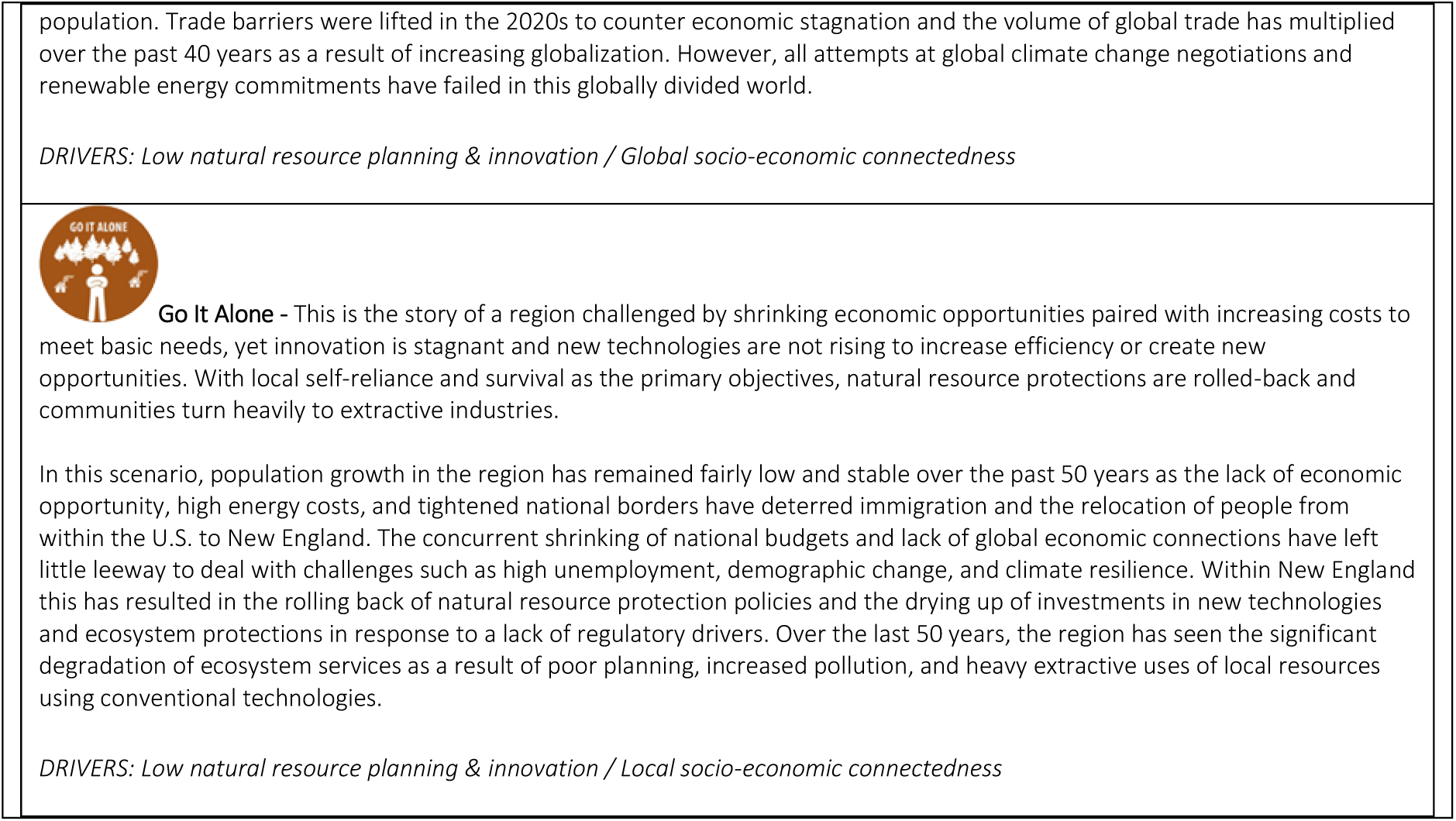
Scenario Narratives

Here our objectives are to: 1) assess the utility and challenges of translating qualitative scenarios into spatial simulations using a cellular LCM; 2) evaluate the outcomes of the scenarios in terms of the differences in the LULC configuration relative to the *Recent Trends* scenario and to each other; 3) compare the fate the landscape in terms of development and conservation within key Impact Areas (i.e., areas that have been identified as being important for conservation, wetland, flood, drinking water, farmland, and or wildlife management) (Figure 2). (4) make the scenarios and simulations available to New England land use stakeholders.

**Figure 2.**
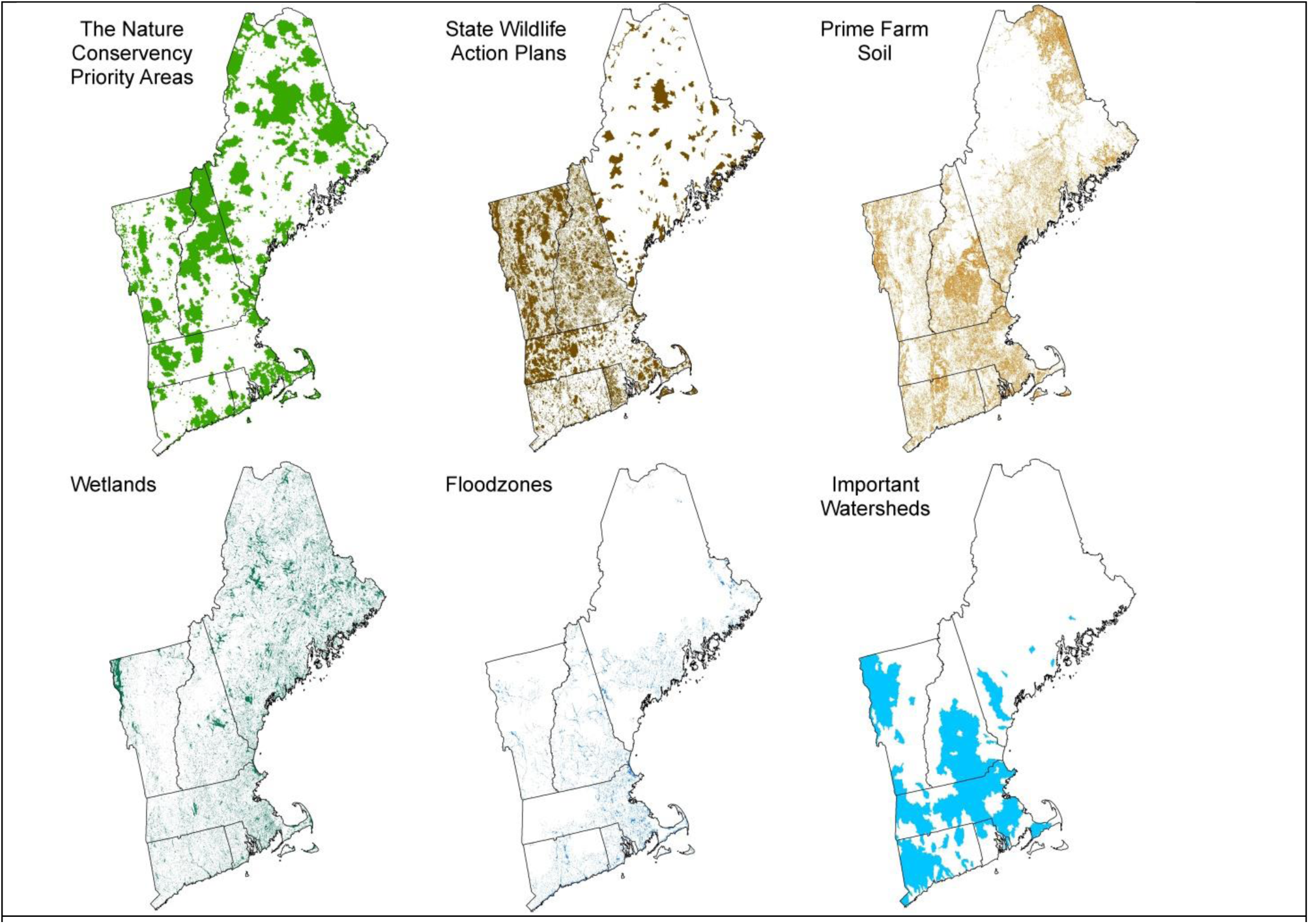
Conservation Priority Areas

## METHODS

### Study Region

New England has a land area of 162,716 km^2^ and includes the six most northeasterly states in the U.S.: Maine (80,068 km^2^), Vermont (23,923 km^2^), New Hampshire (23,247 km^2^), Massachusetts (20,269 km^2^), Connecticut (12,509 km^2^) and Rhode Island (2,700 km^2^) (Figure 1). In 2010, the nominal starting date for the scenarios, 80.1% of the region was forest cover, 7.3% was low density development defined as development with <50% impervious cover, 1.3% was high density development defined as development with >50% impervious cover, and 6.4% was agricultural cover. These estimates were calculated from two sources: (1) the 2010 land cover map produced by Olofsson et al. (Olofsson et al. 2016) applying the Continuous Change Detection and Classification (CCDC) algorithm to Landsat data for all of Massachusetts, New Hampshire, and Rhode Island, 93% of Vermont, 99% of Connecticut, and approximately 33% of Maine and (2) the 2011 National Land Cover Dataset (NLCD), also a Landsat product, for the remainder of New England (Homer et al. 2012). The CCDC and NLCD maps were reclassified to a common legend consisting of: High Density Development, Low Density Development, Forest, Agriculture, Water, and a composite “Other” class that consisted of landcovers such as bare rock and, wetlands which made up less than 5% of the landscape at year 2010 (Appendix I, table 1).

To account for regional variation in the patterns and drivers of land-cover change, we delineated 32 subregions within New England (Figure 1) and independently fit the LCM to the rate and spatial allocation of change within each subregion. The subregions primarily follow U.S. Census Bureau defined Core Base Statistical Areas (CBSA), which represent both Census Metropolitan and Micropolitan statistical areas (www.census.gov; accessed 4/20/2019). CBSAs are delineated to include a core area containing a substantial population nucleus, together with adjacent towns and communities that are integrated with the core in terms of economic and social factors. New England includes 27 CBSAs, however not all of New England is covered by a CBSA. Accordingly, we added five rural areas to fill the gaps, for a total of 32 unique subregions. Among subregions, the Boston-Cambridge-Newton subregion (hereafter “Boston”) is, by far, the most populous; it contains the city of Boston, which is the region’s largest city, and in 2010 accounted for 31% of the region’s total population.

### The simulation framework

We used the Dinamica Environment for Geoprocessing Objects (Dinamica EGO v.2.4.1) to simulate fifty years (2010 to 2060) of LULC change for each scenario, using ten-year time steps. Dinamica EGO is a spatially explicit LCM capable of multi-scale simulations that incorporate spatial feedbacks (Soares-Filho et al. 2002, 2009). The model has several attributes that make it well suited to simulating alternative LULC scenarios. Users prescribe the rate of each potential transition (Figure 3), the ratio of new vs. expansion patches, the mean and variance of new patch sizes, and patch shape complexity. The conditional probability of each transition is developed in relation to a suite of spatial predictor variables. When simulations are intended to project the pattern of LULC change observed in the past, Dinamica EGO employs a weights-of-evidence approach to set the transition probability for every pixel (Soares-Filho et al. 2009). This method is based on a modified form of Bayes theorem of conditional probability; it derives weights such that the effect of each spatial variable on a LULC transition is calculated independently. We used this approach to develop the spatial allocation of land-use to simulate a *Recent Trends* scenario in New England (Thompson et al. 2017) then modified the conditional probabilities to simulate the alternative scenarios (see below).

**Figure 3.**
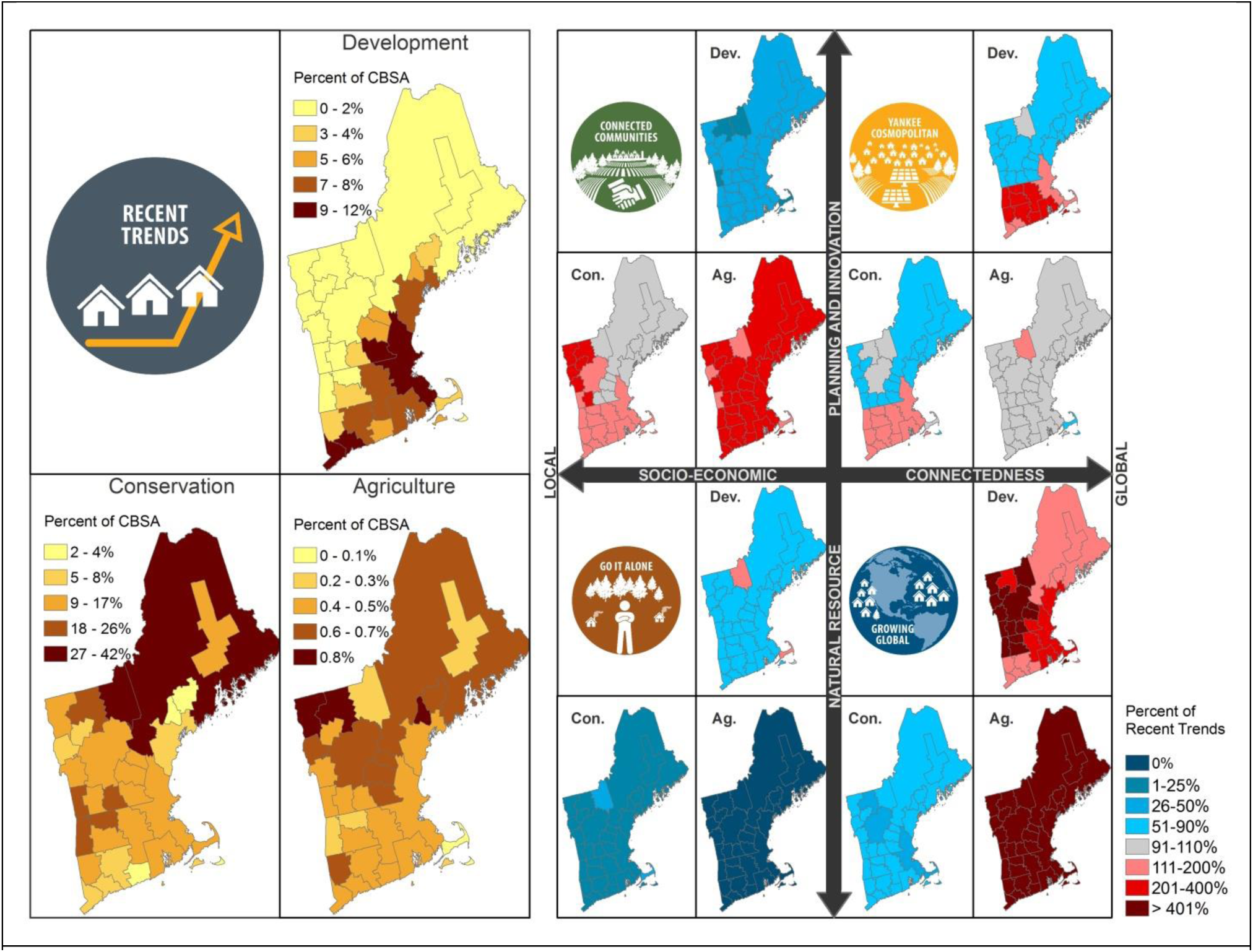
Left: Annual quantity of land use and land cover change with the Recent Trends Scenarios by subregion. Right: Percent change from recent trends for each alternative scenario and land-use landcover change

### Simulating co-designed scenarios

We simulated each of the five LULC change scenarios using Dinamica (Figure 4). The first scenario, the *Recent Trends*, projects the types, rates, and spatial allocation of land cover change and land protection observed during the period spanning 1990 to 2010. Thompson et al. (2017) described the approach for simulating the *Recent Trends* scenario; all LULC transitions in the alternative scenarios were simulated using the same approach. For every LULC transition type, the rate, and allocation observed within each subregion was applied to each time step in the simulation.. For the *Recent Trends* scenario, the transition rate and spatial allocation of the transitions was based on the conversion rate, average patch sizes, ratios of new patch to patch expansion, and patch shape complexity found within the transitions observed in the 1990 to 2010 reference period. The spatial distribution of LULC change was based on observed relationship to eight predictor variables (Table 2). When a subregion could not accommodate a new LULC transition, any remaining unfulfilled transitions were evenly distributed to neighboring subregions. This allowed high development growth subregions like Boston (#7) to spill over into neighboring subregions. The exception to this rule was the island subregions of Nantucket (#28) and Martha’s Vineyard (#3), which were not allowed to spill over since they had no neighboring subregions.

**Table 2.** Driver variables.

| Table 2. Driver variables. |  |  |  |
| --- | --- | --- | --- |
| Variable | Units | Minimum Bin Size | Source |
| Distance to Development | Meters | 100 m | Olofsson et al. 2016 |
| Distance to Cities with population > 30,000 | Meters | 10,000 m | U.S. Department of the Census 1990, 2010 |
| Distance to Roads/Highways | Meters | 100 m | Olofsson et al. 2016 |
| Slope | Degrees | 2° | U.S. Department of the Census 1990, 2010 |
| Land Owner Type | Categorical | NA | Sewall GIS Services 2015<br>http://www.sewall.com/services/geospatial/gis.php |
| Wetlands | Categorical | NA | U.S. Fish and Wildlife Service 2016, Federal Emergency Management Agency 2016, United States Geological Service 2016. |
| Population Density | People per Square Kilometer | 25 ppl/sq. km. | U.S. Department of the Census 1990, 2010. |
| Farm Soil | Categorical | NA | U.S. Department of Agriculture 2016. |

**Figure 4.**
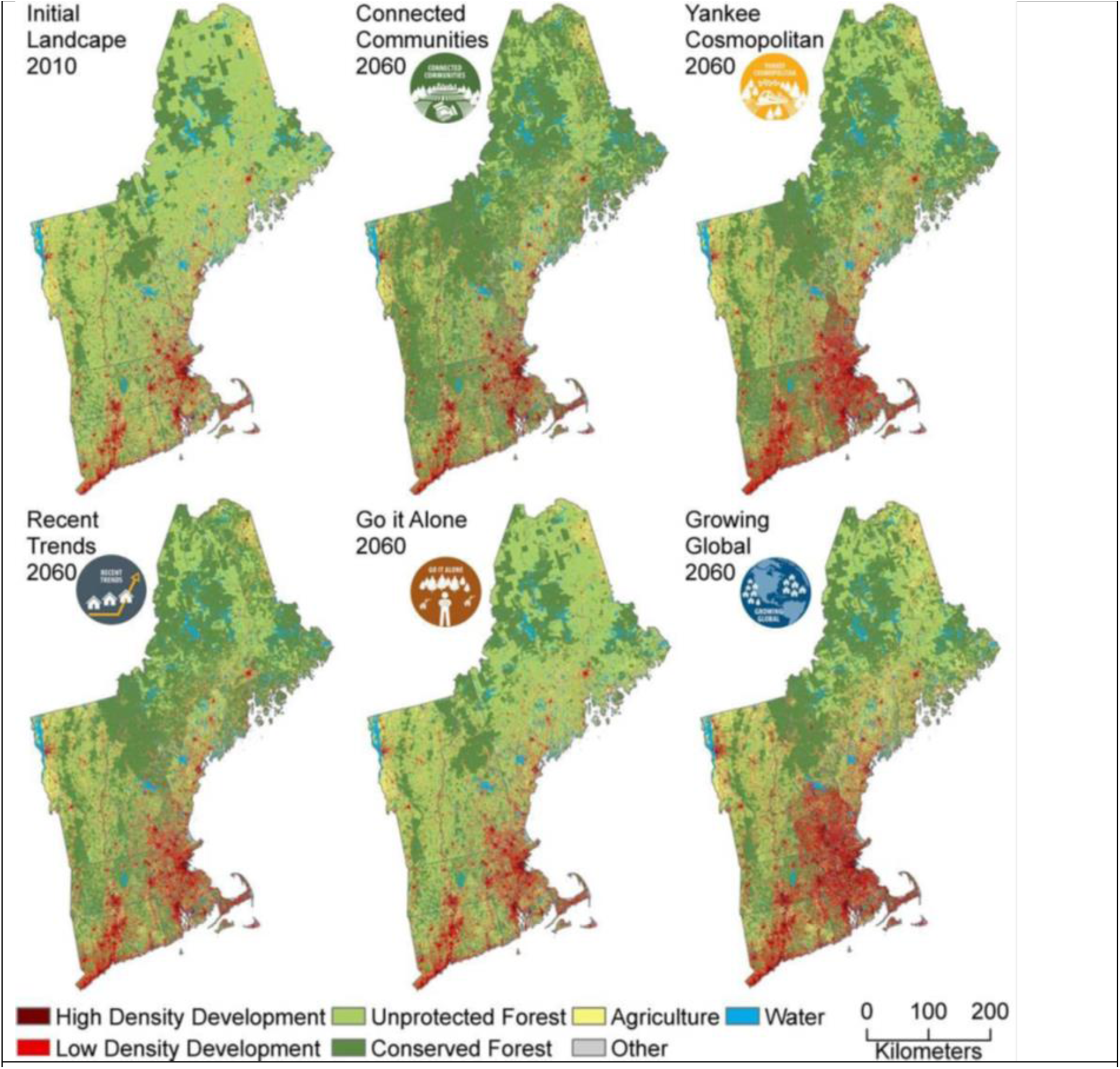
Maps of land cover and land use within New England initial conditions at year 2010, and five alternative scenarios at year 2060.

The four co-designed scenarios have many distinct characteristics of LULC change; they are: *Yankee Cosmopolitan, Connected Communities, Go it Alone*, and *Growing Global* (Box 1). The spatial distribution of each land use in each scenario varied across the landscape and among the scenarios (Figure 4). We used the qualitative descriptions of land-use change provided by the stakeholders in the scenario narratives to develop and propose spatial allocation plans for the land-use transitions in the co-designed scenarios. These spatial allocation plans were presented to the stakeholders in terms of modifications to the baseline weights calculated for the *Recent Trends* scenario. These modifications were then vetted with the stakeholders via webinars and online real-time polling to assess whether they accurately captured their intended deviation from the spatial patterns present in *Recent Trends*. For example, the *Connected Communities* scenario narrative stated that “New settlements tend to occur in planned urban centers”; in response, we suggested that the probability of development be increased as a function of proximity to urban centers and, in a webinar, the stakeholders voted on one of three such modifications that differed in terms of the magnitudes of the adjustment. Table 3 shows the final spatial allocation plans in conjunction with their corresponding quotes from the scenario narratives. The stakeholders assumed that shifts in the LULC change regime would take some time to deviate from the *Recent Trends* rate, so in the first ten-year time step, the rates of LULC change ramp up or down to half of their final target rate (Figure 5).

**Table 3.**
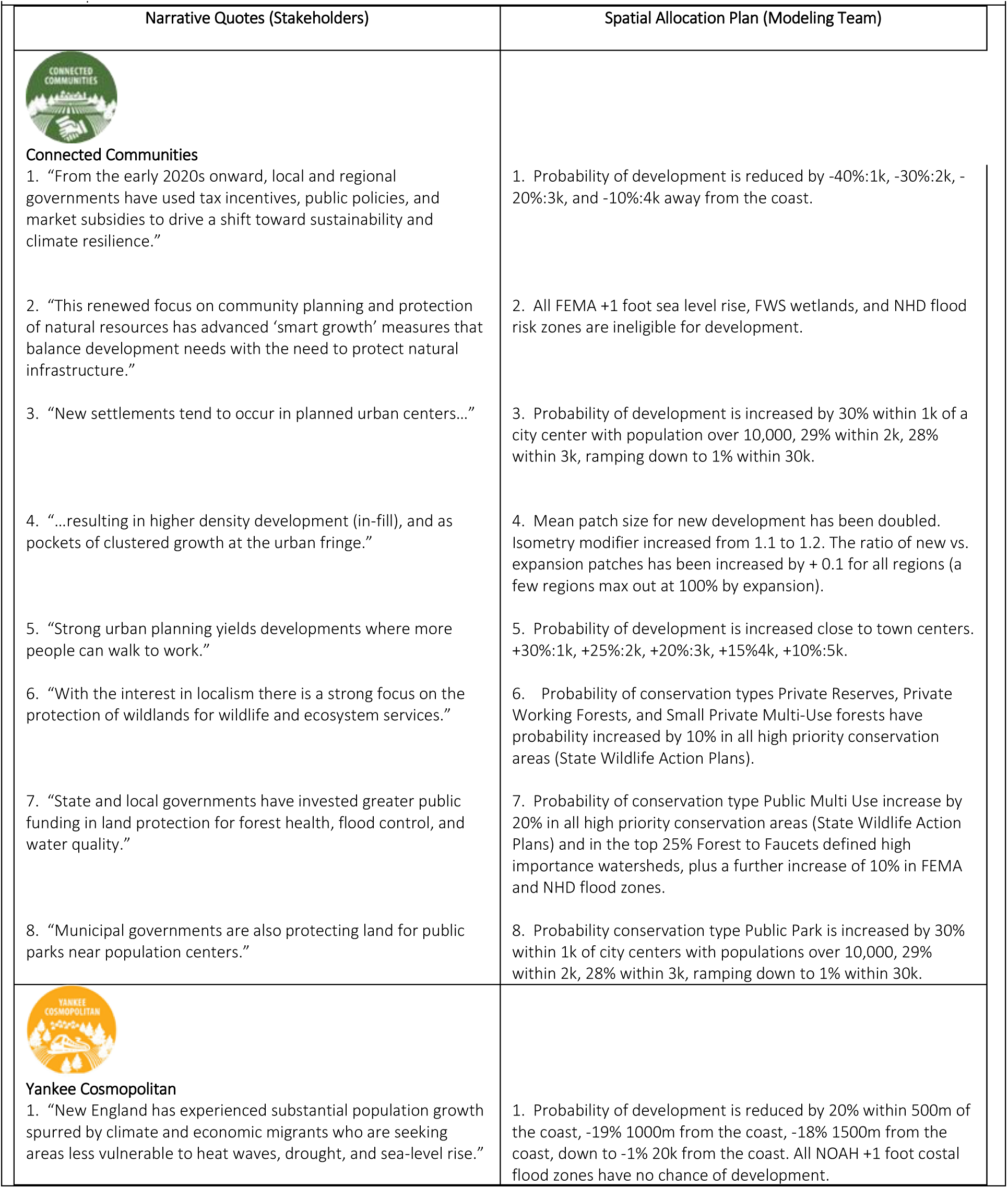
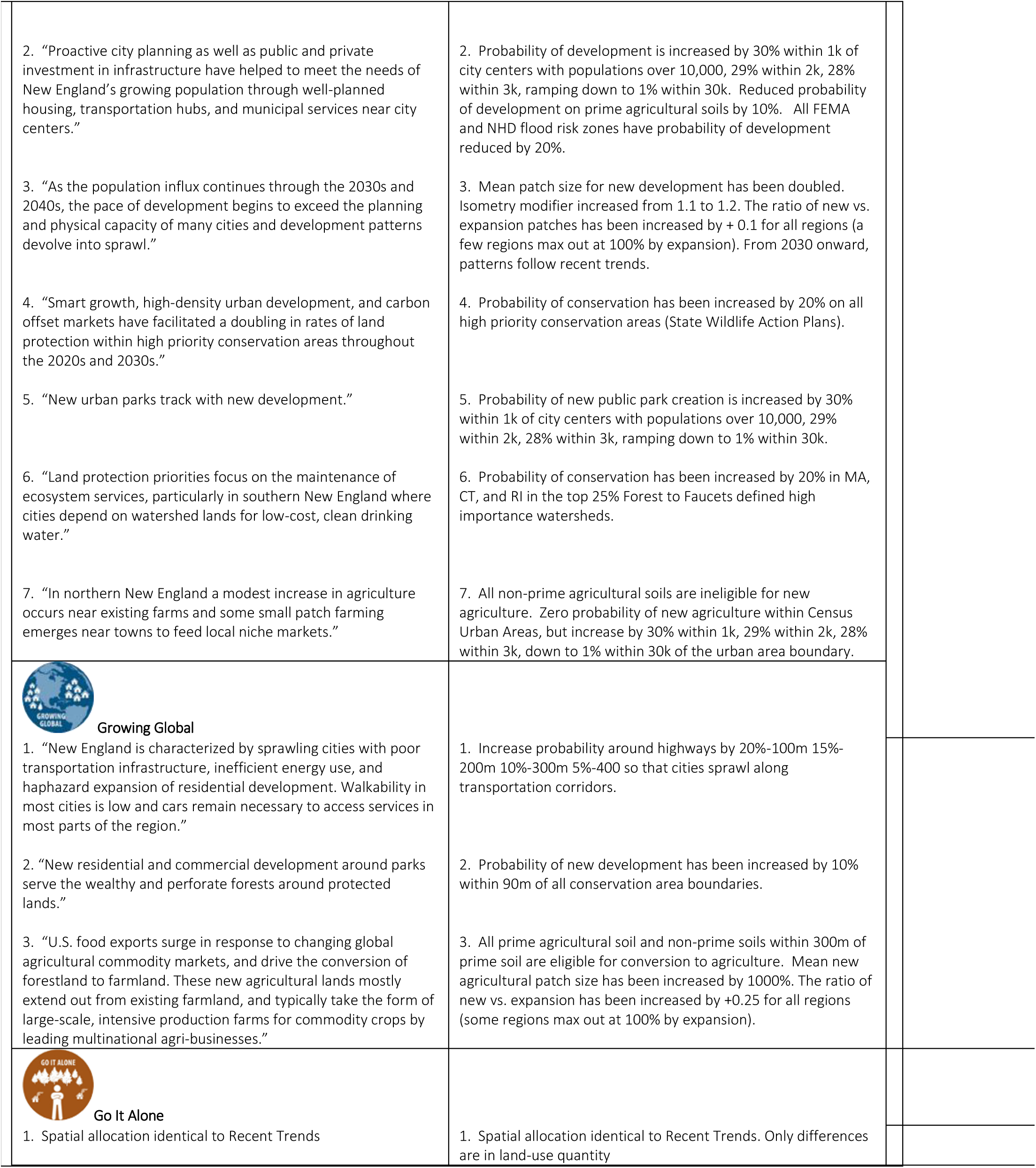
Spatial Allocation Plans

| Table 3. Spatial Allocation Plans |  |
| --- | --- |
| Narrative Quotes (Stakeholders) | Spatial Allocation Plan (Modeling Team) |
| **Connected Communities**<br>1. "From the early 2020s onward, local and regional governments have used tax incentives, public policies, and market subsidies to drive a shift toward sustainability and climate resilience."<br>2. "This renewed focus on community planning and protection of natural resources has advanced 'smart growth' measures that balance development needs with the need to protect natural infrastructure."<br>3. "New settlements tend to occur in planned urban centers..."<br>4. "...resulting in higher density development (in-fill), and as pockets of clustered growth at the urban fringe."<br>5. "Strong urban planning yields developments where more people can walk to work."<br>6. "With the interest in localism there is a strong focus on the protection of wildlands for wildlife and ecosystem services."<br>7. "State and local governments have invested greater public funding in land protection for forest health, flood control, and water quality."<br>8. "Municipal governments are also protecting land for public parks near population centers." | 1. Probability of development is reduced by -40%:1k, -30%:2k, -20%:3k, and -10%:4k away from the coast.<br>2. All FEMA +1 foot sea level rise, FWS wetlands, and NHD flood risk zones are ineligible for development.<br>3. Probability of development is increased by 30% within 1k of a city center with population over 10,000, 29% within 2k, 28% within 3k, ramping down to 1% within 30k.<br>4. Mean patch size for new development has been doubled. Isometry modifier increased from 1.1 to 1.2. The ratio of new vs. expansion patches has been increased by + 0.1 for all regions (a few regions max out at 100% by expansion).<br>5. Probability of development is increased close to town centers. +30%:1k, +25%:2k, +20%:3k, +15%:4k, +10%:5k.<br>6. Probability of conservation types Private Reserves, Private Working Forests, and Small Private Multi-Use forests have probability increased by 10% in all high priority conservation areas (State Wildlife Action Plans).<br>7. Probability of conservation type Public Multi Use increase by 20% in all high priority conservation areas (State Wildlife Action Plans) and in the top 25% Forest to Faucets defined high importance watersheds, plus a further increase of 10% in FEMA and NHD flood zones.<br>8. Probability conservation type Public Park is increased by 30% within 1k of city centers with populations over 10,000, 29% within 2k, 28% within 3k, ramping down to 1% within 30k. |
| **Yankee Cosmopolitan**<br>1. "New England has experienced substantial population growth spurred by climate and economic migrants who are seeking areas less vulnerable to heat waves, drought, and sea-level rise." | 1. Probability of development is reduced by 20% within 500m of the coast, -19% 1000m from the coast, -18% 1500m from the coast, down to -1% 20k from the coast. All NOAA +1 foot coastal flood zones have no chance of development. |
| 2. "Proactive city planning as well as public and private investment in infrastructure have helped to meet the needs of New England's growing population through well-planned housing, transportation hubs, and municipal services near city centers."<br>3. "As the population influx continues through the 2030s and 2040s, the pace of development begins to exceed the planning and physical capacity of many cities and development patterns devolve into sprawl."<br>4. "Smart growth, high-density urban development, and carbon offset markets have facilitated a doubling in rates of land protection within high priority conservation areas throughout the 2020s and 2030s."<br>5. "New urban parks track with new development."<br>6. "Land protection priorities focus on the maintenance of ecosystem services, particularly in southern New England where cities depend on watershed lands for low-cost, clean drinking water."<br>7. "In northern New England a modest increase in agriculture occurs near existing farms and some small patch farming emerges near towns to feed local niche markets." | 2. Probability of development is increased by 30% within 1k of city centers with populations over 10,000, 29% within 2k, 28% within 3k, ramping down to 1% within 30k. Reduced probability of development on prime agricultural soils by 10%. All FEMA and NHD flood risk zones have probability of development reduced by 20%.<br>3. Mean patch size for new development has been doubled. Isometry modifier increased from 1.1 to 1.2. The ratio of new vs. expansion patches has been increased by + 0.1 for all regions (a few regions max out at 100% by expansion). From 2030 onward, patterns follow recent trends.<br>4. Probability of conservation has been increased by 20% on all high priority conservation areas (State Wildlife Action Plans).<br>5. Probability of new public park creation is increased by 30% within 1k of city centers with populations over 10,000, 29% within 2k, 28% within 3k, ramping down to 1% within 30k.<br>6. Probability of conservation has been increased by 20% in MA, CT, and RI in the top 25% Forest to Faucets defined high importance watersheds.<br>7. All non-prime agricultural soils are ineligible for new agriculture. Zero probability of new agriculture within Census Urban Areas, but increase by 30% within 1k, 29% within 2k, 28% within 3k, down to 1% within 30k of the urban area boundary. |
| **Growing Global**<br>1. "New England is characterized by sprawling cities with poor transportation infrastructure, inefficient energy use, and haphazard expansion of residential development. Walkability in most cities is low and cars remain necessary to access services in most parts of the region."<br>2. "New residential and commercial development around parks serve the wealthy and perforate forests around protected lands."<br>3. "U.S. food exports surge in response to changing global agricultural commodity markets, and drive the conversion of forestland to farmland. These new agricultural lands mostly extend out from existing farmland, and typically take the form of large-scale, intensive production farms for commodity crops by leading multinational agri-businesses." | 1. Increase probability around highways by 20%-100m 15%-200m 10%-300m 5%-400 so that cities sprawl along transportation corridors.<br>2. Probability of new development has been increased by 10% within 90m of all conservation area boundaries.<br>3. All prime agricultural soil and non-prime soils within 300m of prime soil are eligible for conversion to agriculture. Mean new agricultural patch size has been increased by 1000%. The ratio of new vs. expansion has been increased by +0.25 for all regions (some regions max out at 100% by expansion). |
| **Go It Alone**<br>1. Spatial allocation identical to Recent Trends | 1. Spatial allocation identical to Recent Trends. Only differences are in land-use quantity |

**Figure 5.**
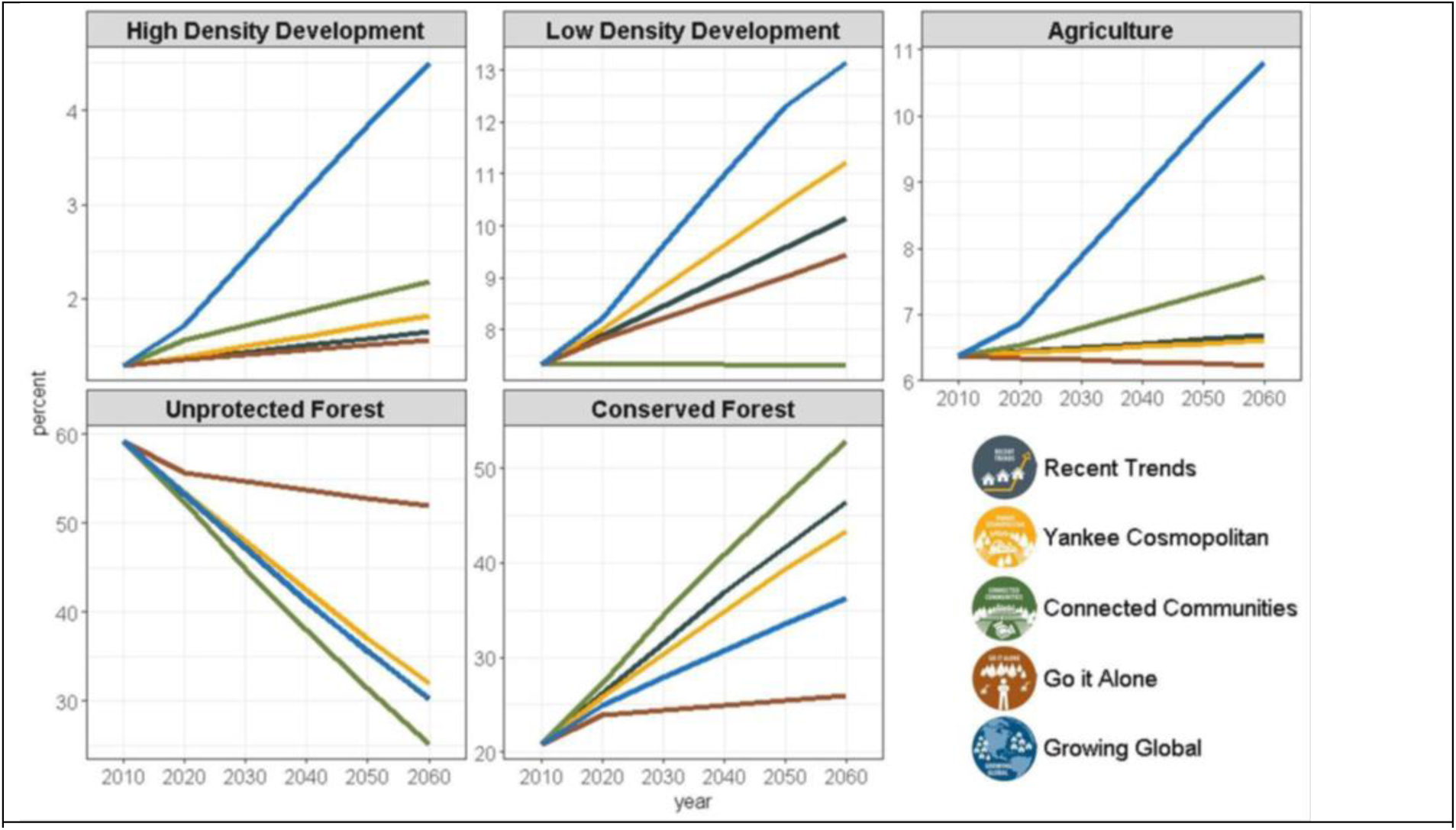
Changes in land cover within New England over time for each LULC class and scenario. Note varying Y-axes.

### Scenario Impacts on Conservation Priorities

To explore the impacts of the scenarios, we estimated the impacts of simulated LULC change on forests within each scenario on the following seven key Impact Areas. We selected these areas because they serve as reasonable proxies for a range of values and conditions that are important to stakeholders (McBride et al. 2019) and have been mapped previously within New England.

i. *Core Forests*, were delineated as forested areas that are >30 meters from a non-forest land cover at the start of the simulation (i.e., in 2010).
ii. *Flood Zones*, were defined where Federal Emergency Management Agency FEMA Flood Zones with 1% annual probability of flooding (Zones A, AE, AH, AO, and VE) (Federal Emergency Management Agency 2017). Note that not all subregions have FEMA-defined Flood zones.
iii. *Surface Drinking Water*, were defined as the 25% highest scoring watersheds classified by the US Forest Service Forest to Faucets report (Weidner and Todd 2011). Watersheds were ranked based on the importance of their surface water quality in relation to the human demand on that water supply.
iv. *Wildlife Habitats*, were delineated using State Wildlife Action Plans (SWAP) (New Hampshire Fish and Game Department 2012, Maine Dept. of Inland Fisheries and Wildlife 2015, Massachsetts Division of Fisheries and Wiliflife 2015, Rhode Island Department of Environmental Management Division of Fish and Wildlife 2015, State of Connecticut Department of Energy and Enviornmental Protection 2015, Vermont Fish & Wildlife Department 2015). We accounted for state level variation in wildlife conservation priorities and for the variable proportion of land given priority status by focusing on the top tiers of each state’s Wildlife Habitat priorities as high-value wildlife conservation assets and then standardized the scores by scaling them relative to the mean score for all land in each state. Therefore, wildlife habitat values greater than 1.0 indicate areas with better than average wildlife value.
v. *TNC Priority Conservation Areas,* was delineated based on The Nature Conservancy’s Priority Conservation Areas. These areas aim to represent the full distribution and diversity of native species, natural communities, and ecosystems such that a conserving these areas will ensure the long-term survival of all native life and natural communities, not just threatened species and communities.
vi. *Wetlands,* were defined as wetlands classified by the National Wetlands Inventory Wetlands (U S Fish and Wildlife Service 2012).
vii. *Prime Farmlands,* were identified using the Farmland Class from the Gridded Soil Survey Geographic (gSSURGO) Database (SSURGO Soil Survey Staff 2011). We merged the Farmland Classes: farmland of statewide importance, all areas are prime farmland, farmland of unique importance, and farmland of local importance into one “Prime Farmlands” classification.

Impact Areas were assessed based on the amount of land available for conversion to either development or conservation at the start of the simulations in 2010. Areas already developed or conserved in 2010 were considered unavailable and were thus not assessed. Additionally, areas within delineated Impact Areas that were ineligible for a transition based on our model rules (e.g. non-forest covers such as agriculture, water and other) were not considered.

### Developing outreach tool

We used the scenarios and simulation products to develop an online interactive mapping tool to portray the interaction between land use choices and land use outcomes in New England and support efforts by community groups and conservation groups to explore how they might adapt their LULC plans and conservation priorities to ensure that they are robust under an uncertain future. The tool, the NELF Explorer (www.newenglandlandscapes.org) was built by FernLeaf Interactive and the National Environmental Modeling and Analysis Center (NEMAC) at the University of North Carolina Asheville. The NELF Explorer was built using the simulation outputs in consultation with user perspectives, via a project launch visioning session plus three cycles of prototyping and user-review. Users can use the NELF Explorer to navigate among five scenarios (Recent Trends, Go it Alone, Connected Communities, Yankee Cosmopolitan, and Growing Global) and visualize how each scenario influences land use and ecosystem services at 5 time-points (2010, 2020, 2030, 2040, 2050, 2060), across all six New England states, at multiple scales including state, county, town, and watershed. The NELF Explorer displays maps with land use color coded (High Density Development, Low Density Development, Unprotected Forest, Conserved Forest, Agriculture, and Water). Graphs show the number of acres in each type of land use for each scenario at the six time-points. Also, the outcomes of scenario comparisons in 2060 for Impact Areas of Flood Zones, Surface Drinking Water, Wildlife Habitats, Priority Conservation Areas, Wetlands, Prime Farmland, and Core Forests are described within the tool. The tool is static; the underlying data and calculations were completed in advance via the simulation process. Therefore, the NELF Explorer is a conduit for accessing pre-computed data and visualizations.

## RESULTS

### Recent Trends

The *Recent Trends* scenario assumes a continuation of the LULC changes observed between 1990 and 2010. The rate of LULC change is constant throughout the scenario: New development covers 97 km^2^ per year; new agriculture covers 16 km^2^ per year; and new land protection covers in 835 km^2^ per year. At year 2060 (after simulating 50 years of LULC change), developed land increased by 37% (from 14,098 to 19,265 km^2^); there was little change (< 5%) in agricultural land cover (10,409 to 10,908 km^2^). The largest LULC change was to protected land, which increased by 123% (from 35,300 to 78,500 km^2^).

Throughout the fifty-year simulation, the rate of land protection in the *Recent Trends* scenario was more than eight times greater than the rate of development. Because Impact Areas are not evenly distributed throughout New England, the spatial distribution of land protection in the *Recent Trends* scenario was most effective for securing protection in Impact Areas that are concentrated in the north, such as Core Forest, where 48% was protected and only 3% developed and *TNC Priority Conservation Areas* where 49% was protected and only 4% developed. Impact Areas that are concentrated in the south, such as with the Important Watersheds for Drinking water only 28% was Protected and11% was developed. In addition, the impact of LULC change on other conservation priorities was driven by local patterns observed in the historical data. For example, wetlands have regulatory protection (included in our model) and thus have a low probability of development. Indeed, despite being common throughout the region, 45% of forested wetland areas were protected while just 0.7% were developed (note that non-forested wetlands were protected from any transition).

### Yankee Cosmopolitan

The *Yankee Cosmopolitan* scenario envisions a future New England that is a global hub of activity, with commensurate changes to land use. The population is growing much faster than *Recent Trends*, but, at the same time, natural resource planning and innovation are a priority. To accommodate population growth spurred by climate and economic migrants, development occurred at a rate 40% greater than *Recent Trends* (136 km^2^ per year). Global food supply chains required minimal agriculture expansion, which was maintained at 16 km^2^ per year (the same as *Recent Trends*). The rate of new land protection was reduced in the north and increased in the south, relative to *Recent Trends*. Overall, across the region, the rate of land protection in this scenario was 736km^2^ per year, 12% lower than *Recent Trends.*

*Yankee Cosmopolitan* includes several modifications to the spatial allocation of LULC change in *Recent Trends,* which were intended to minimize development within areas desirable for protection. However, the large (40%) increase in the rate of development often overwhelmed modifications to the spatial allocation rules. For example, the spatial allocation plan for *Yankee Cosmopolitan* included a reduced probability of new development within flood zones (Table 3); nonetheless, forest loss within flood zones by year 2060 was 86% higher than in *Recent Trends*. Reduced development probability in flood zones was only effective in rural subregions, where there was less development pressure. In urbanizing subregions, where development rates were highest even low probability sites were eventually developed. Similarly, the spatial allocation plan for this scenario increased the probability of land protection within wildlife habitat areas; however, the increased rate of development had a greater influence. Overall, while there was a small increase in protected land within wildlife habitat areas, there was also a 49% increase in developed areas, as compared to *Recent Trends*. Other modifications to the spatial allocation were more effective. For example, this scenario envisioned more urban parks thus the spatial allocation plan increased the probability of new protected lands within two km of city centers, which resulted in a 75% increase in protected areas within two km of city centers, compared to the *Recent Trends* scenario. In addition, concentrating development around city centers resulted in a similar amount of core forest to the *Recent Trends*, despite accommodating 40% more development.

### Connected Communities

The *Connected Communities* scenario envisions a future characterized by local socio-economic connectedness and high natural resource planning and innovation. Population growth slowed and became more compact and, as a result, the rate of new development was just 25% of the rate in the *Recent Trends*—24 km^2^ per year. Local agriculture expanded to meet the need for local food and forests were converted to new agricultural land at a rate of 41 km^2^ per year, more than 248% of the rate of forests to agriculture simulated in *Recent Trends*. This scenario also included a strong focus on land protection for wildlife and ecosystem services; the rate of new land protection was 1045 km^2^ year.

Consistent with this scenario’s emphasis on natural resource conservation and planning, the spatial allocation of LULC change in the *Connected Communities* scenario included a lower probability of development and increased probability of land protection within flood zones, wildlife habitat areas and important drinking water watersheds. These modifications, combined with a lower overall rate of new development, resulted in: a 77% decrease in the amount of development in flood zones by 2060; an 80% decrease in the amount of development in wildlife habitat areas; and 71% increase in land protection in drinking water important watersheds. Indeed, the *Connected Communities* scenario had the greatest increase in the amount of protected land within the Impact Areas across all the scenarios. The scenario narrative emphasized compact development and the simulation of the scenario had the greatest proportion of new development was within 10 km of cities among all scenario (XX% more development within 10km of cities than *Recent Trends)*. As part of this scenario’s emphasis on climate change adaptation, the proportion of development within 5-km of the coast (where sea-level rise is a concern) was significantly less than *Recent Trends*.

### Go It Alone

The *Go It Alone* scenario envisions a future with low natural resource planning and innovation and local socio-economic connectedness. New England has shrinking economic opportunities and communities turn heavily to extractive industries. Rates of land development slowed to 75km^2^ per year, which was a 25% reduction from *Recent Trends*. Where development continued, it was characterized by unplanned residential housing that perforates the landscape. There was no new agriculture cover. Land protection tapered off dramatically early in the scenario and by 2060 there was 80% less new protected land than in the *Recent Trends* scenario.

While the rates are much lower, the spatial allocation of LULC change in *Go It Alone* followed the patterns developed for the *Recent Trends* Scenario. Less new development resulted in proportionately less forest loss within Impact Areas, including 25% less priority wildlife habitat loss and 31% less development on flood plains. Relatedly, the large reduction in the rate of land protection resulted in *Go It Alone* having the lowest level of conservation within Impact Areas among the five scenarios.

### Growing Global

The *Growing Global* scenario envisions and landscape undergoing massive changes. Migration into New England drives up the population. Local planning efforts have failed to keep pace with development. Economic and social connectivity is globalized while natural resource planning and innovation is low. Compared to the *Recent Trends* scenario, *Growing Global* resulted in an 182% increase in the rate of new development, a 900% increase in the rate of new agriculture, and a reduction of 40% in the rate of new land protection.

In this scenario, the total amount of developed land in New England more than doubled (from 14,090 to 28,880 km^2^) by 2060. Boston grew to a sprawling mega city the size of modern day Tokyo, Japan. Rapid and largely unregulated development resulted in the greatest increase in development within Impact Areas among all scenarios. For example, the *Growing Global* scenario did not include any spatial modifier to decrease the probability of development in flood zones or other Impact Areas. As a result, by 2060, this scenario developed 275% more flood zones compared to the *Recent Trends* scenario. There were similarly high (+275%) increases in development within high priority wildlife habitats. More than twice as much land near the coast (<10km) was developed, as compared to the *Recent Trends*.

## DISCUSSION and CONCLUSIONS

Our process for translating co-designed qualitative scenarios into quantitative simulations of LULC change yielded divergent representations of the future New England landscape. The simulations differ markedly in terms of the amount of LULC change and the spatial pattern of change. Indeed, among scenarios there is a fivefold difference in the amount of high-density development, and a twofold difference in the amount of protected land. While all the scenarios represent distinct storylines resulting in discrete manifestations of those stories, the *Growing Global* scenario stands out for having, by far, the greatest amount of change. By year 2060, *Growing Global* envisions that urban expansion around Boston will sprawl to an area covering more than 10,000 km2, larger in size than Tokyo, Japan. On one hand, this is such a drastic change that it may seem implausible to stakeholders and thereby undermine the utility of the scenario. On the other hand, the simulation is faithful to the stakeholders’ storyline, which envisions New England as a destination for millions of migrants fleeing the growing impacts of climate change elsewhere (National Climate Assessment 2018). Specifically, the stakeholders describe: “sprawling cities with poor transportation infrastructure, inefficient energy use, and haphazard expansion of residential development.” The plausibility of this scenario is supported anecdotally by events such as Hurricane Maria, which, in 2017, displaced as many as 500,000 people from the island of Puerto Rico to the mainland U.S. (Pew Research Center 2018). Given that a single storm can cause such large changes to settlement patterns, it will be important to consider the consequences of scenarios, such as *Growing Global* which push our assumptions about how the past can or cannot shape the future. Overall, the simulated scenarios bound a wide range of future possibilities for the New England landscape and, as such, have high potential for broadening the perspectives of planners, counteracting a general tendency toward ‘narrow-thinking’ when planning for an uncertain future (Soll et al. 2014).

Our simulations effectively captured the land-use dynamics and features described in the scenario storylines. Each specific modification to *Recent Trends* is annotated within the qualitative scenario descriptions so that our stakeholders can see how their vision for each scenario was incorporated into the simulation. By identifying specific quotes that referenced differences in land-use patterns, then translating them into explicit rules for the spatial allocation of simulated LULC change (Table 3), we were able to capture the intentions of the stakeholders in ways that had substantive and readily attributable impacts on the simulated landscape. For example, simulated development surrounding the area of Keene, New Hampshire (subregion 24) in *Go it Alone* and *Yankee Cosmopolitan* both have the same rate of development but different spatial allocation of that development (Figure 6). The *Yankee Cosmopolitan* narrative described: “Proactive city planning as well as public and private investment in infrastructure have helped to meet the needs of New England’s growing population through well-planned housing, transportation hubs, and municipal services near city centers.” Thus, a spatial modifier was implemented in this scenario to concentrate development close to city centers while protecting farm soils and limiting development in flood zones (Table 3). Overall this approach represents an effective and transparent method for bridging the gap between non-technical stakeholders who developed the scenarios and the technical experts who simulated them (Mallampalli et al. 2017). We are hopeful that this clear translation of the scenarios to the simulations bolsters the legitimacy and salience of the participatory scenario process (*sensu* Cash et al 2002) and results in greater use by the stakeholders and decision-makers.

**Figure 6.**
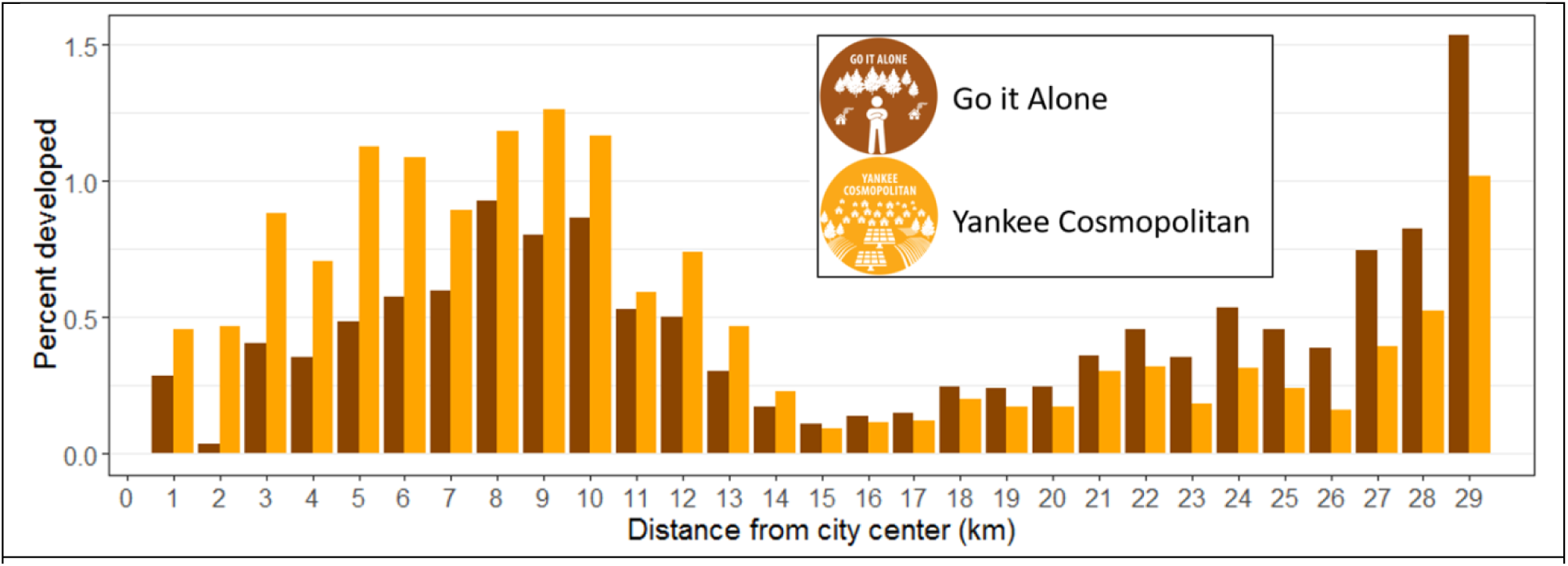
Spatial Allocation Example. Distance to Keene, NH city center. Two scenarios with same amount of development but different spatial allocation.

These simulations reveal much about the potential impacts of future land use on conservation priorities. In general, the amount of projected LULC change affected the Impact Areas more than the differences in their spatial allocation. For example, the *Yankee Cosmopolitan* scenario has several spatial allocation rules designed to mitigate the impacts to conservation goals, including: reduced probability of new development within flood zones and increased probability of land protection within wildlife habitat areas. In comparison, the Go It Alone scenario has no modifications to the spatial allocation rules. However, *Yankee Cosmopolitan* has **87%** more development than *Go it Alone*. So despite substantial efforts to mitigate the impacts of development, the *Yankee Cosmopolitan* scenario resulted in more development in every category of Impact Area than *Go it Alone*. This pattern is consistent across all scenarios and Impact Areas, insomuch as the rank order of development within each impact area matched the rank order of the amount of development, despite strong differences in the spatial allocation patterns (Figure 7).

**Figure 7.**
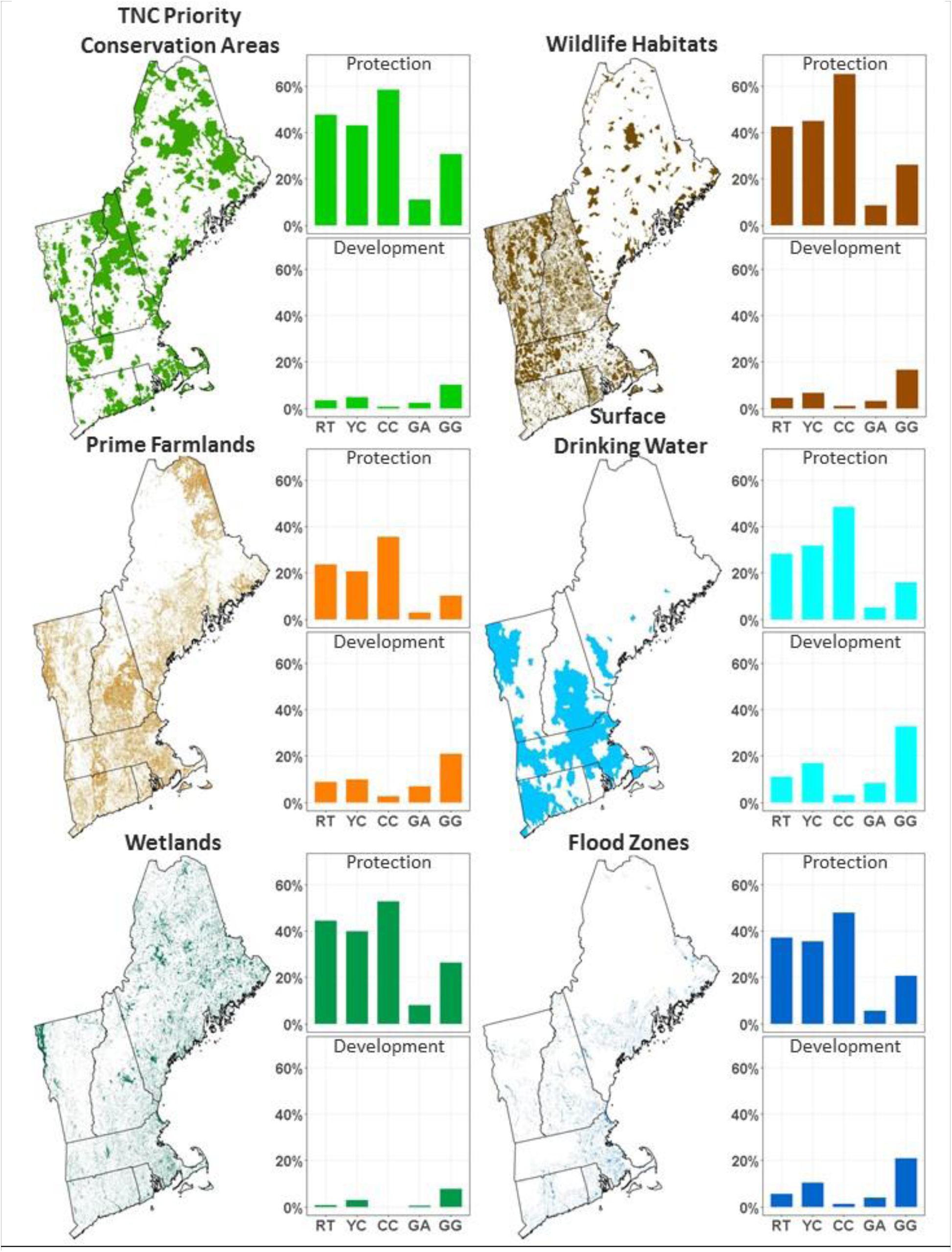
Impact Areas. Inset bar charts represent the percent of each conservation priority area that was developed (bar left of zero), and conserved (bar right of zero) for each scenario at year 2060.

The simulated land-cover scenarios were designed to meet multiple goals. One key goal was to create simulated land-cover scenarios that catalyze new research which to understand and advance sustainable land-use trajectories. In addition to the analyses presented here, our hope is that the scenarios will serve as a common platform that brings researchers together to examine the consequences of changing land use. To that end, all the spatial layers (i.e., GIS maps) from this project are available on Data Basin^1^, an open-source spatial data repository. Indeed, researchers from around the region have begun to use the simulation outputs within other landscape models to explore how these scenarios affect various ecosystem services and landscape outcomes.

Our final goal was to make the scenarios and simulations available to New England land use stakeholders to promote future scenario thinking at the community scale and provide a spatial analysis tool for evaluating risks to specific lands and conservation goals from the local to regional scale. For this community of users, we developed the New England Landscape Futures (NELF) Explorer^2^. The tool was designed via a user-engagement process to meet the needs of diverse stakeholders, including conservationists, planners, developers, government leaders, and citizens who want to explore possible land-use futures in specific areas. The NELF Explorer was launched in March 2019. We are currently tracking use of the tool and collaborating with NELF Explorer users to document use cases. Potential uses of the NELF Explorer include understanding the future of the land through local scenario planning, conservation and development planning, and community engagement/education.

## ACKNOWLEDGEMENTS

This research was supported in part by National Science Foundation funded to the Harvard Forest Long Term Ecological Research Program (Grant No. NSF-DEB 12-37491) and the Scenarios Society and Solutions Research Coordination Network (Grant No. NSF-DEB-13-38809). We thank the commitment and energy offered by the participants of the scenario development process, and Jeff Hicks, Jim Fox, Karin Rogers, and their team at FernLeaf Interactive and the National Environmental Modeling and Analysis Center (NEMAC) at the University of North Carolina Asheville, for developing the NELF Explorer.

**Appendix I Table 1.**

| Reclassification Scheme |  |  |  |  |
| --- | --- | --- | --- | --- |
| THIS STUDY | CCDC Class | CCDC Class Description | NLCD 2001/2011 Class | NLCD 2001/2011 Class Description |
| High Density Developed | Commercial/Industrial | Area of urban development; impervious surface area target 80-100% | Developed High Intensity | Highly developed areas where people reside or work in high numbers. Examples include apartment complexes, row houses and commercial /industrial. Impervious surfaces account for 80% to 100% of the total cover. |
|  | High Density Residential | Area of residential urban development with some vegetation; impervious surface area target 50-80% | Developed, Medium Intensity | Areas with a mixture of constructed materials and vegetation. Impervious surfaces account for 50% to 79% of the total cover. These areas most commonly include single-family housing units. |
| Low Density Developed | Low Density Residential | Area of residential urban development with significant vegetation; impervious surface area target 0-50% | Developed, Low Intensity | Areas with a mixture of constructed materials and vegetation. Impervious surfaces account for 20% to 49% percent of total cover. These areas most commonly include single-family housing units. |
|  |  |  | Developed, Open Space | Areas with a mixture of some constructed materials, but mostly vegetation in the form of lawn grasses. Impervious surfaces account for less than 20% of total cover. These areas most commonly include large-lot single-family housing units, parks, golf courses, and vegetation planted in developed settings for recreation, erosion control, or aesthetic purposes. |
| Agriculture | Agriculture | Non-woody cultivated plants; includes cereal and broadleaf crops | Pasture/Hay | Areas of grasses, legumes, or grass-legume mixtures planted for livestock grazing or the production of seed or hay crops, typically on a perennial cycle. Pasture/hay vegetation accounts for greater than 20% of total vegetation. |
|  |  |  | Cultivated Crops | Areas used for the production of annual crops, such as corn, soybeans, vegetables, tobacco, and cotton, and also perennial woody crops such as orchards and vineyards. Crop vegetation accounts for greater than 20% of total vegetation. This class also includes all land being actively tilled. |
| Forest | Mixed Forest | Forested land with at least 40% tree canopy cover comprising no more than 80% of either evergreen needleleaf or deciduous broadleaf cover | Mixed Forest | Areas dominated by trees generally greater than 5 meters tall, and greater than 20% of total vegetation cover. Neither deciduous nor evergreen species are greater than 75% of total tree cover. |
|  | Deciduous Broadleaf Forest | Forested land with at least 40% tree canopy cover comprising more than 80% deciduous broadleaf cover | Deciduous Forest | Areas dominated by trees generally greater than 5 meters tall, and greater than 20% of total vegetation cover. More than 75% of the tree species shed foliage simultaneously in response to seasonal change. |
|  | Evergreen Needleleaf Forest | Forested land with at least 40% tree canopy cover comprising more than 80% evergreen needleleaf cover | Evergreen Forest | Areas dominated by trees generally greater than 5 meters tall, and greater than 20% of total vegetation cover. More than 75% of the tree species maintain their leaves all year. Canopy is never without green foliage. |
|  | Woody Wetland | Additional class of wetland that tries to separate wetlands with considerable biomass from mainly herbaceous wetlands | Woody Wetlands | Areas where forest or shrubland vegetation accounts for greater than 20% of vegetative cover and the soil or substrate is periodically saturated with or covered with water. |
|  |  |  | Shrub/Scrub | Areas dominated by shrubs; less than 5 meters tall with shrub canopy typically greater than 20% of total vegetation. This class includes true shrubs, young trees in an early successional stage or trees stunted from environmental conditions. |
| Other | Wetland | Vegetated land (woody and non-woody) with inundation from high water table; includes swamps, salt and freshwater marshes and tidal rivers/mudflats | Emergent Herbaceous Wetlands | Areas where perennial herbaceous vegetation accounts for greater than 80% of vegetative cover and the soil or substrate is periodically saturated with or covered with water. |
|  | Herbaceous / Grassland | Non-woody naturally occurring or slightly managed plants; includes pastures | Barren Land (Rock/Sand/Clay) | Areas of bedrock, desert pavement, scarps, talus, slides, volcanic material, glacial debris, sand dunes, strip mines, gravel pits and other accumulations of earthen material. Generally, vegetation accounts for less than 15% of total cover. |
|  | Bare | Non-vegetated land comprised of above 60% rock, sand, or soil |  |  |
| Water | Water | Lakes, ponds, rivers, and ocean | Open Water | Areas of open water, generally with less than 25% cover of vegetation or soil. |

## Footnotes

1 https://databasin.org/groups/26ceb6c7ece64b0d9872e118bae80d41

2 www.newenglandlandscapes.org

